# Feed-forward metabotropic signaling by Cav1 Ca^2+^ channels supports pacemaking in pedunculopontine cholinergic neurons

**DOI:** 10.1101/2023.08.05.552108

**Authors:** C. Tubert, E. Zampese, T. Pancani, T. Tkatch, D. J. Surmeier

**Affiliations:** Department of Neuroscience, Feinberg School of Medicine, Northwestern University, Chicago, IL 60611, USA; Universidad de Buenos Aires - CONICET. Instituto de Fisiología y Biofísica Bernardo Houssay (IFIBIO Houssay), Facultad de Medicina, Departamento de Ciencias Fisiológicas. Grupo de Neurociencia de Sistemas. Buenos Aires, Argentina; Max Planck Florida Institute for Neuroscience, 1 Max Planck Way, Jupiter, Florida 33458, USA; Aligning Science Across Parkinson’s (ASAP) Collaborative Research Network, Chevy Chase, MD 20815 USA

## Abstract

Like a handful of other neuronal types in the brain, cholinergic neurons (CNs) in the pedunculopontine nucleus (PPN) are lost in the course of Parkinson’s disease (PD). Why this is the case is unknown. One neuronal trait implicated in PD selective neuronal vulnerability is the engagement of feed-forward stimulation of mitochondrial oxidative phosphorylation (OXPHOS) to meet high bioenergetic demand, leading to sustained oxidant stress and ultimately degeneration. The extent to which this trait is shared by PPN CNs is unresolved. To address this question, a combination of molecular and physiological approaches were used. These studies revealed that PPN CNs are autonomous pacemakers with modest spike-associated cytosolic Ca^2+^ transients. These Ca^2+^ transients were attributable in part to the opening of high-threshold Cav1.2 Ca^2+^ channels, but not Cav1.3 channels. Nevertheless, Cav1.2 channel signaling through endoplasmic reticulum ryanodine receptors stimulated mitochondrial OXPHOS to help maintain cytosolic adenosine triphosphate (ATP) levels necessary for pacemaking. Inhibition of Cav1.2 channels led to recruitment of ATP-sensitive K^+^ channels and slowing of pacemaking. Cav1.2 channel-mediated stimulation of mitochondria increased oxidant stress. Thus, PPN CNs have a distinctive physiological phenotype that shares some, but not all, of the features of other neurons that are selectively vulnerable in PD.

## Introduction

PD is the second most common neurodegenerative disorder (Poewe et al., 2017). The cardinal motor symptoms of PD – bradykinesia, rigidity and tremor – can be ascribed to the degeneration of dopaminergic neurons in the substantia nigra pars compacta (SNc). However, PD has both motor and non-motor features that cannot be readily attributed to the loss of these neurons. Indeed, PD Lewy pathology is found in many brain regions with the most prominent sites being in the medulla (Surmeier, Obeso, et al., 2017). One of the medullary nuclei affected relatively early in PD is the pedunculopontine nucleus (PPN) (Hirsch et al., 1987; Zweig et al., 1989; Shinotoh et al., 1999; Tubert et al., 2019). The neurons found in the PPN can be divided into cholinergic, glutamatergic and GABAergic subsets (Clements et al., 1991; Clements & Grant, 1990; Ford et al., 1995; Lavoie & Parent, 1994). The PPN subset of cholinergic neurons (CNs) have long, highly branched axons that stretch rostrally to the di- and telencephalon, as well as caudally into the brainstem (Semba & Fibiger, 1992; Lavoie & Paren, 1994; Takakusaki et al., 1996; Mena-Segovia et al., 2008; Dautan et al., 2014; 2016). These PPN CNs are more vulnerable in PD than neighboring glutamatergic and GABAergic neurons (Hirsch et al., 1987).

The determinants of this selective vulnerability are unknown. Study of other types of neuron that are vulnerable in PD has identified several potential cellular risk factors [Surmeier et al. 2017]. SNc dopaminergic neurons, for example, have long, highly branched axons, autonomous pacemaking and basal mitochondrial oxidant stress. This basal stress appears to be a consequence of feed-forward stimulation of mitochondrial oxidative phosphorylation (OXPHOS) (Zampese et al., 2022). The feed-forward signaling cascade couples activity-dependent plasma membrane Ca^2+^ channels with a Cav1 pore-forming subunit to endoplasmic reticulum (ER) ryanodine receptors (RYRs) and Ca^2+^ entry into ER-docked mitochondria. Ca^2+^ entry into mitochondria stimulates OXPHOS and adenosine triphosphate production. In SNc dopaminergic neurons, engagement of this feed-forward bioenergetic control mechanism is necessary to sustain spiking over extended periods of time.

It is unclear whether this neuronal phenotype is common to other neurons at-risk in PD, particularly those utilizing acetylcholine as a neurotransmitter. Dorsal motor nucleus of the vagus (DMV) cholinergic neurons, which manifest Lewy pathology in PD patients but are not lost with disease progression, exhibit some of the features of SNc dopaminergic neurons (Goldberg et al., 2012; Surmeier, Obeso, et al., 2017). In contrast to DMV cholinergic neurons, PPN CNs not only manifest Lewy pathology, they degenerate (Hirsch et al., 1987). As noted above, PPN CNs have long, highly branched axons, like SNc dopaminergic neurons, but it is unclear whether they have the same physiological properties. In particular, it’s unclear whether they are autonomous pacemakers and engage feed-forward control of mitochondrial OXPHOS. Our goal was to fill this gap. Using a combination of electrophysiological, optical and genetic approaches, these studies revealed that PPN CNs share many physiological traits with SNc dopaminergic neurons: autonomous pacemaking, feed-forward stimulation of mitochondrial OXPHOS, and basal mitochondrial oxidant stress. However, there were several differences. One was the reliance upon Ca^2+^ channels with a Cav1.2 pore-forming subunit, rather than one with a Cav1.3 subunit; another was the robust engagement of K^+^ channels when cytosolic ATP concentrations fell, leading to a suppression of pacemaking.

## Results

### Autonomous pacemaking in PPN CNs was accompanied by a modest cytosolic Ca^2+^ tranisents

To identify PPN CNs in ex vivo brain slices, mice expressing Cre recombinase under control of the choline acetyltransferase (ChAT) promoter (ChAT-Cre) were crossed with Ai14 mice, in which the expression of the red fluorescent protein tdTomato was regulated by Cre recombinase (Tubert et al., 2016). To validate this approach in PPN, tissue sections from these mice were examined to determine the relationship between tdTomato fluorescence and immunoreactivity for ChAT. As expected, there was excellent alignment of the two in the PPN region (Fig. 1A, B).

**Figure 1:**
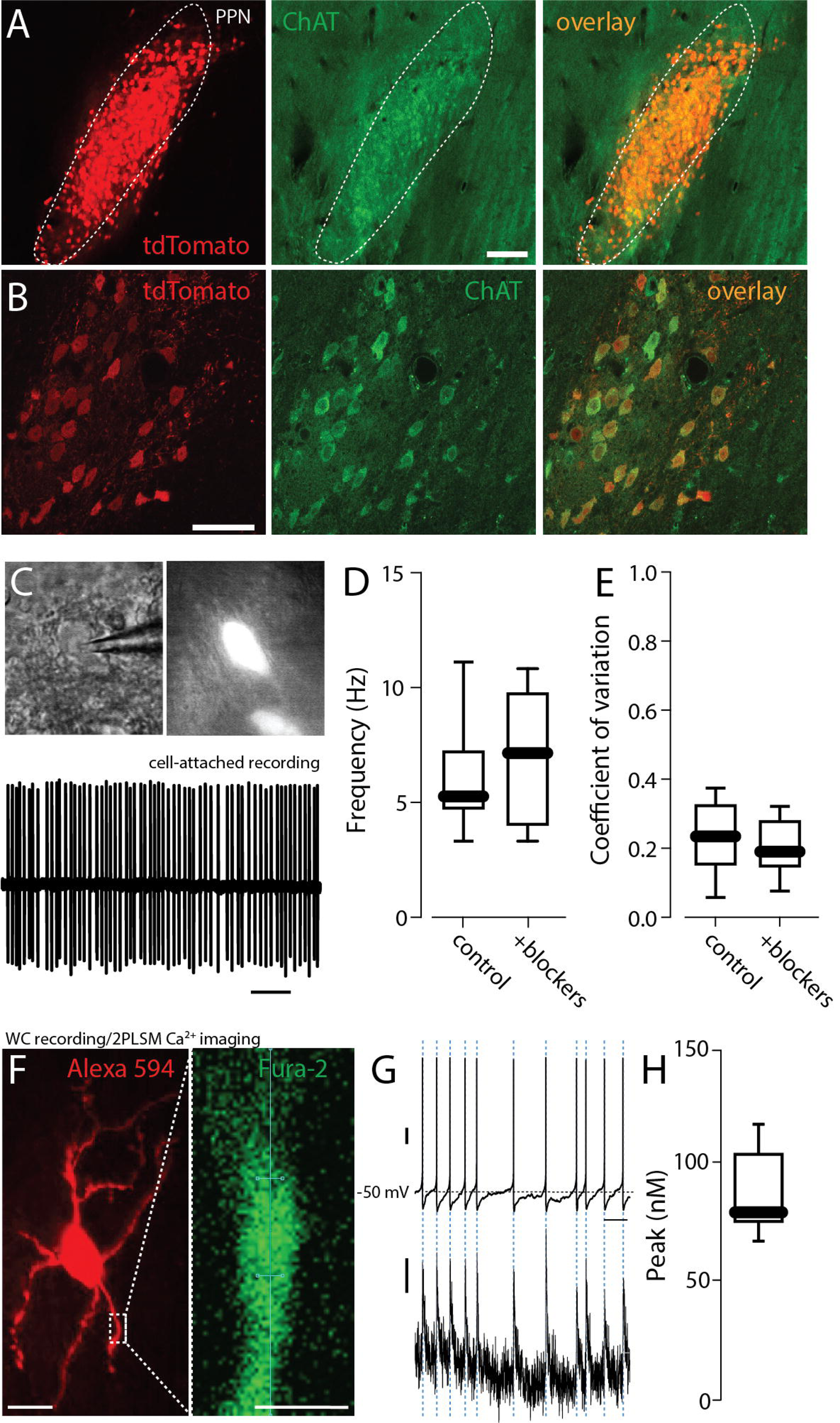
Autonomous pacemaking in PPN CNs was accompanied by a modest cytosolic Ca^2+^ transients. **A.** Representative micrograph of a parasagital section of the PPN from a ChAT-Cre/Ai14 mouse brain showing intrinsic tdTomato fluorescence (red) colocalized with choline acetyltransferase immunolabeling (ChAT, green). Bar: 100 um. **B.** Higher magnification of the PPN showing intrinsic tdTomato fluorescence colocalized with choline acetyltransferase immunolabeling. Bar: 50 um. **C.** Top: Representative micrograph of a PPN-ChAT from a ChAT-Cre/Ai14 mouse under differential interference contrast (DIC) and epifluorescent illumination. Bottom: Representative cell-attached recordings of a PPN CNs in the presence of picrotoxin (100 mM) and CNQX (50 mM). **D-E.** Firing frequency (**D**) and coefficient of variation (**E**) of PPN CNs before and after addition of synaptic blockers picrotoxin and CNQX (Wilcoxon, NS, n=7 in each group). **F.** Representative micrograph of a PPN-CN filled with Alexa 594 (red, top, 50 uM) and the calcium indicator Fura-2 (100 mM). Bar: 20 um. Magnification of a dendrite (right) of a PPN CN filled with Fura-2. Bar: 5 um. **G.** Representative whole-cell voltage recording of PPN CNs spontaneous activity (top; vertical bar: 10 mV) and dendrite calcium oscillations (bottom; vertical bar: 10 nM) recorded during the spontaneous activity. Horizontal bar: 1 s. **H.** Calcium oscillations peak during the spontaneous activity of PPN CNs (n=9). Box plots indicate first and third quartiles, thick center lines represent medians, and whiskers indicate the range.

To determine whether the spontaneous activity of PPN CNs was autonomously generated, fluorescent neurons in the PPN region were monitored using cell-attached patch methods in *ex vivo*, parasagittal brain slices (Fig. 1C). Bath application of antagonists for ionotropic glutamate (CNQX, 50 uM) and GABA (picrotoxin, 100 uM) receptors had no significant effect on the rate or regularity of PPN CN spiking (Fig. 1D,E). Thus, the spontaneous spiking in PPN CNs was autonomously generated.

To determine if spiking was accompanied by fluctuations in intracellular Ca^2+^ concentration, visually-identified PPN CNs were subjected to whole cell recording using electrodes filled with the ratiometric Ca^2+^ indicator Fura-2 (100 uM) (Figure 1F) and fluorescence measured with 2-photon laser-scanning microscopy (2PLSM) (Fig. 1F). Cytosolic Ca^2+^ transients were aligned with somatic spikes (Fig. 1G). Estimates of the peak Ca^2+^ concentration of the transients in proximal dendrites yielded a median value near 75 nM (Fig. 1G), considerably below values observed in SNc dopaminergic neurons (Guzman et al., 2018; Zampese et al., 2022).

In SNc dopaminergic neurons, both Cav1.2 and Cav1.3 Ca^2+^ channels contribute to the cellular mechanisms underlying pacemaking and spiking driven by synaptic input (Guzman et al., 2009; Shin et al., 2022; Zampese et al., 2022). To determine if these channels were playing a similar role in PPN CNs, an adenoassociated virus (AAV) carrying a Cre-dependent expression plasmid for RiboTag (Sanz et al., 2009) and a green fluorescent protein (GFP) reporter was stereotaxically injected into the PPN of ChAT-Cre mice (Fig. 2A,B). The messenger ribonucleic acid (mRNA) harvested from these mice was then subjected to RNASeq analysis (Sanz et al., 2009). Surprisingly, this analysis suggested that the gene coding for the Cav1.3 subunit (*CACNA1D*) was expressed at very low levels, but that the gene coding for the Ca_v_1.2 subunit (*CACNA1C)* was robustly expressed (data not shown). This inference was subsequently confirmed with quantitative reverse transcriptase polymerase chain reaction (qPCR) analysis of RiboTag harvested mRNA (Fig. 2C).

**Figure 2:**
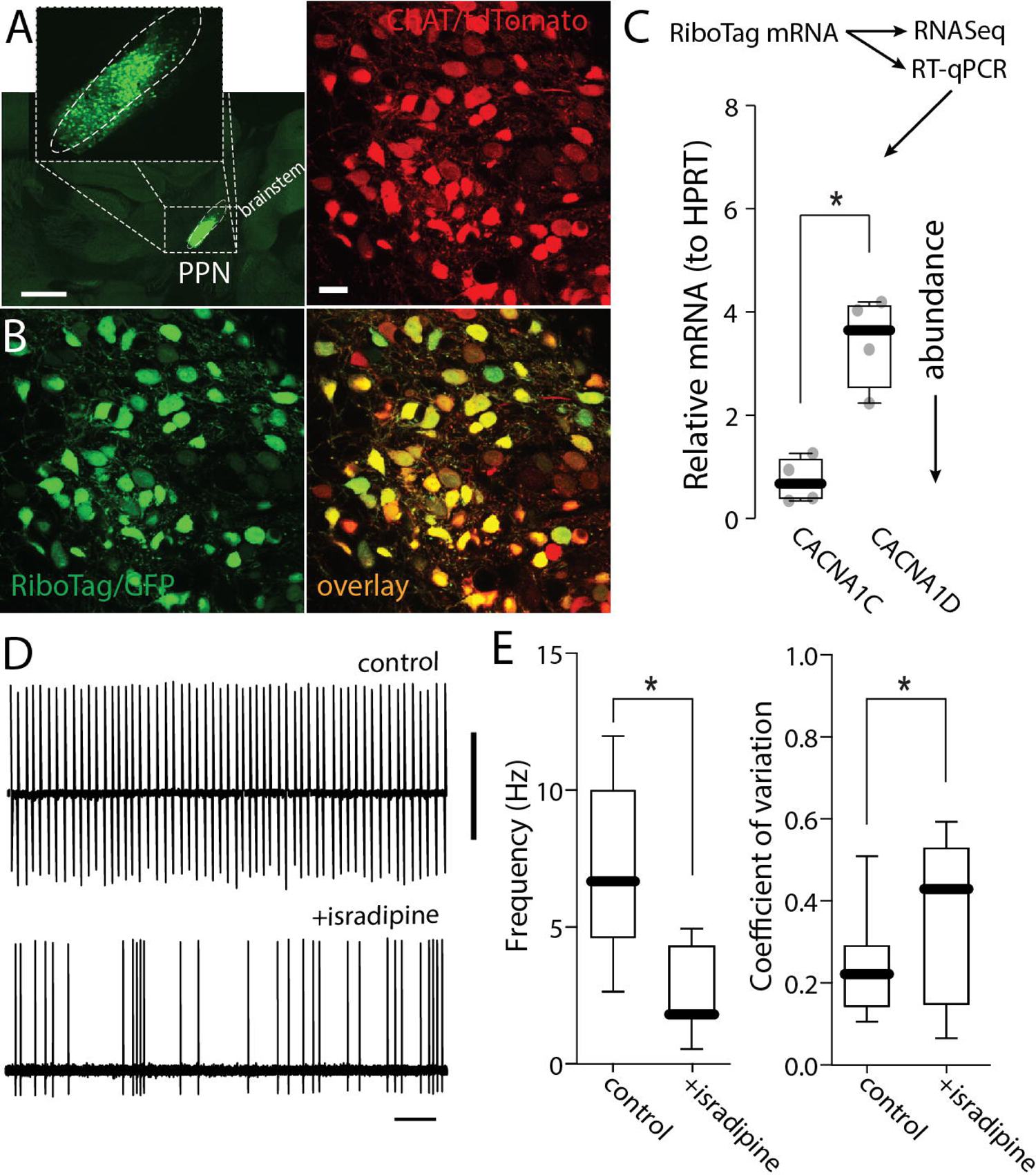
Calcium channels expression in PPN-ChNs. A. Representative micrograph of a parasagital section from a ChAT-Cre/Ai14 mouse brain showing Ribotag-GFP expression (green) in the PPN. Bar: 1000 μm. **B.** Representative micrograph from a ChAT-Cre/Ai14 mouse showing Ribotag-GFP expression (green) in PPN CNs (red). Bar: 20 μm. **C.** Quantification of RNA abundance of Ca_v_1.2 (CACNA1C) and Ca_v_1.3 (CACNA1D) (t-test; *p=0.0071; n=4 mice). **D.** Representative cell-attached recordings of PPN CNs without or with chronic isradipine (500 nM, 1 hr pre-incubation). Vertical bar: 5 mV; horizontal bar: 1 s. **E.** Firing frequency and coefficient of variation of PPN CNs with or without isradipine (Frequency: Mann-Whitney, *p=0.0030; coefficient of variation: Mann-Whitney, *p=0.0016; n=11 in each group). Box plots indicate first and third quartiles, thick center lines represent medians, and whiskers indicate the range.

To explore the potential functional consequences of the Cav1 subunit expression profile, pharmacological experiments were performed. Cav1.3 and Cav1.2 Ca^2+^ channels differ in the voltage-dependence of channel activation: Cav1.3 channels open at sub-spike threshold membrane potentials and Cav1.2 channels only open at supra-threshold potentials after the spike has been triggered. Thus, Cav1.3 channels contribute to the depolarization necessary to drive pacemaking. On the other hand, being ‘high threshold’ Cav1.2 channels are not suited to directly participate in the ionic mechanisms driving the depolarization necessary to maintain pacemaking. As a consequence, it was surprising to find that bath application of isradipine (1 µM), a negative allosteric modulator of Cav1 channels, significantly slowed pacemaking and decreased its regularity (Fig. 2D-E).

### Cav1 channels trigger ER Ca^2+^ release and Ca^2+^ entry into mitochondria

Given that the effects of isradipine on PPN CNs are largely attributable to inhibition of Cav1.2 channels, why did isradipine slow pacemaking? One way in which Cav1.2 channels could have affected pacemaking is by helping to maintain cytosolic ATP levels that are necessary for ionic homeostasis and suppression of K-ATP channels (Ashcroft, 2005). In SNc dopaminergic neurons, Cav1 Ca^2+^ channels stimulate mitochondrial ATP production by triggering ER Ca^2+^ release through RYRs; Ca^2+^ released from the ER then enters mitochondria to enhance OXPHOS (Zampese et al., 2022). As a first step toward determining whether a similar mechanism was engaged by PPN CNs (Fig. 3A), the sensitivity of mitochondrial matrix Ca^2+^ content to inhibition of plasma membrane Cav1 Ca^2+^ channels was assessed. To perform this assessment, the PPN of ChAT-Cre/Ai14 mice was injected with an AAV carrying a Cre-dependent expression construct for a mitochondrially targeted Ca^2+^ sensor (mitoGCaMP6) (Fig. 3B-E). Several weeks after AAV injection, mice were sacrificed, and ex vivo brain slices were studied using 2PLSM. As in SNc dopaminergic neurons, pharmacological inhibition of plasma membrane Cav1 Ca^2+^ channels with isradipine (1 µM) or inhibition of RYRs with 1,10-diheptyl-4,40-bipyridinium dibromide (DHBP, 100 µM) significantly lowered mitochondrial matrix Ca^2+^ content in PPN CNs (Fig. 3F,G). As expected, DHBP inhibition, was more effective than inhibition of Cav1 channels alone in reducing mitochondrial matrix Ca^2+^ content during autonomous pacemaking.

**Figure 3:**
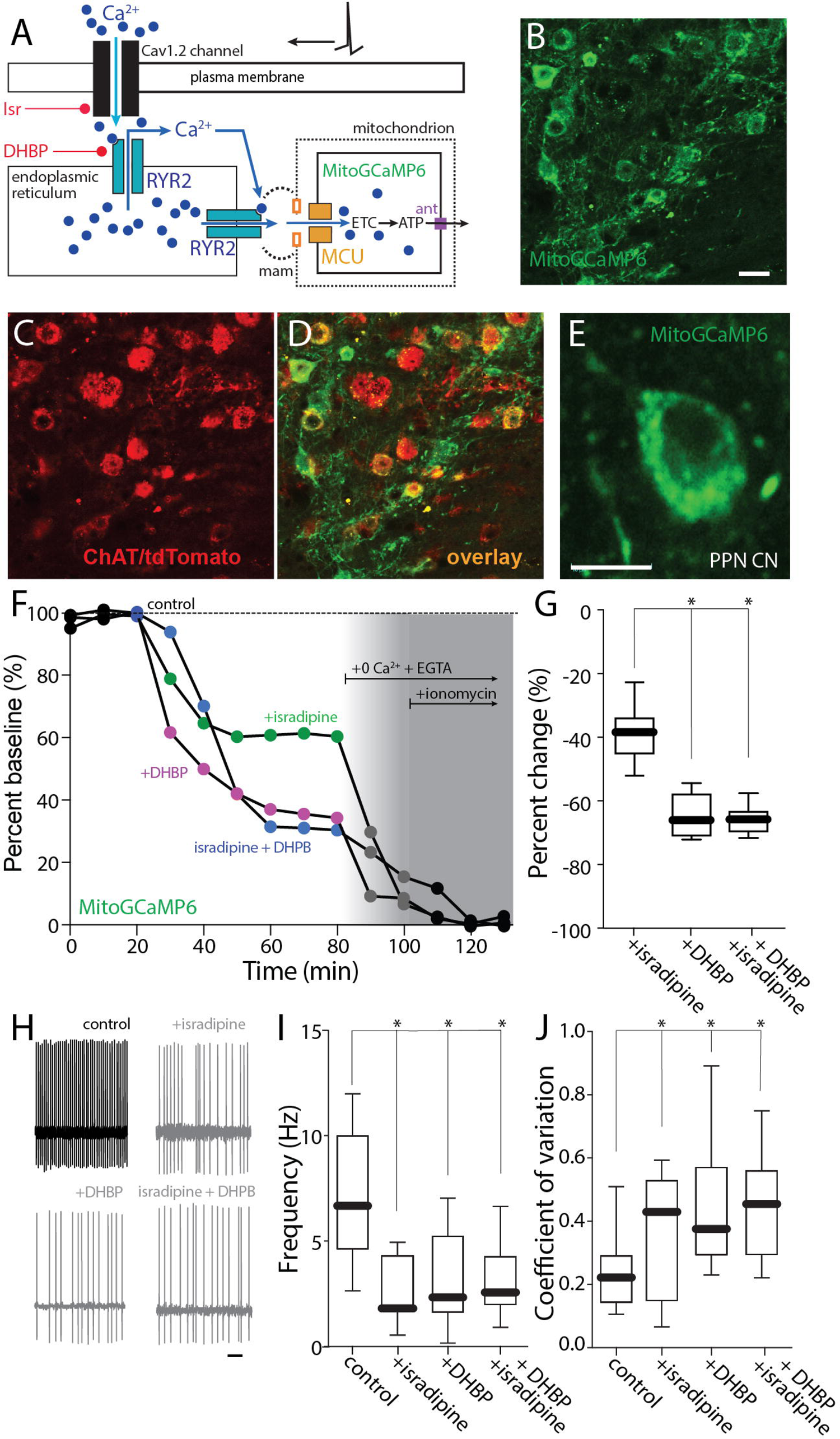
Ca_v_1 channels trigger ER Ca^2+^ release and Ca^2+^ entry into mitochondria. **A.** Cartoon illustrating the effect of Ca^2+^ entry through Cav1.2 channels on RYR Ca^2+^ release. **B-D.** Representative micrograph from a ChAT-Cre/Ai14 mouse showing mitoGCaMP6 expression (**B**) in choline acetyltransferase immunolabeled neurons (ChAT, **C**) and the overlay of both markers (**D**) Bar: 20 μm. **E.** Representative micrograph of a recorded PPN CN expressing mitoGCaMP6. Bar: 10 μm. **F.** Representative example of the percentage of mitochondrial Ca^2+^ in a PPN CN before and after addition of isradipine (500 nM), DHBP or both, 0 Ca^2+^ + EGTA and 0 Ca^2+^ + EGTA + ionomycin, all in physiological glucose concentration (3.5 mM) aCSF. **G.** Percentage of change of mitochondrial Ca^2+^ from aCSF for isradipine, DHBP and both (Kruskal-Wallis, Interaction: *p=0.0001; Dunn’s multiple comparision, Isradipine-DHBP: *p=0.0050; Isradipine-Isradipine+DHBP: *p=0.0043; n=7 in each group). **H.** Representative cell-attached recordings of PPN-ChNs without or with chronic isradipine (500 nM, 1 hr pre-incubation), chronic DHBP (100 μM, 1 hr pre-incubation), or both, in the presence of picrotoxin (100 mM) and CNQX (50 mM) in a low-glucose (3.5 mM) ACSF. Bar: 1 s. **I-J.** Firing frequency (**I**) and coefficient of variation (**J**) of PPN-ChNs with or without isradipine, DHBP, or both (**I:** Non-parametric one-way ANOVA, interaction *p=0.0040; Dunn’s multiple comparision: control-Isradipine: *p=0.0058; control-DHBP: *p=0.0385; control-Isradipine+DHBP: *p=0.0273; Isradipine-DHBP: NS; Isradipine-Isradipine+DHBP: NS; DHBP-Isradipine+DHBP: NS. **J:** Non-parametric one-way ANOVA, interaction *p=0.0033; Dunn’s multiple comparision: control-Isradipine: *p=0.0426; control-DHBP: *p=0.0150; control-Isradipine+DHBP: *p=0.0065; Isradipine-DHBP: NS; Isradipine-Isradipine+DHBP: NS; DHBP-Isradipine+DHBP: NS; n=10-11 in each group). Box plots indicate first and third quartiles, thick center lines represent medians, and whiskers indicate the range.

If the effect of isradipine on pacemaking was mediated by control of mitochondrial matrix Ca^2+^ content, then blocking RYRs should mimic the effects of isradipine. To test this hypothesis, PPN CNs were monitored using cell-attached recording in ex vivo brain slices (as described above). As predicted, blockade of RYRs with DHBP (100 µM) mimicked the effect of isradipine – slowing the pacemaking rate and increasing its irregularity (Fig. 3H-J). Moreover, DHBP occluded the effect of isradipine on pacemaking rate and regularity.

In contrast to commonly used protocols for ex vivo slice recording, these experiments were performed in an external solution containing physiological concentrations of glucose (3.5 mM). In this condition, the contribution of glycolysis to cytosolic ATP levels will be less than when glucose is elevated. To determine how extracellular glucose concentration shaped the effects of isradipine (Fig. 4A), the ex vivo experiments were repeated in the presence of elevated glucose concentration (25 mM). In this condition, the effect of isradipine and DHBP on mitochondrial Ca^2+^ content was not altered (Fig. 4B-D). However, in the presence of high glucose, isradipine and DHBP failed to significantly alter spiking rate or regularity (Fig. 4E-G).

**Figure 4:**
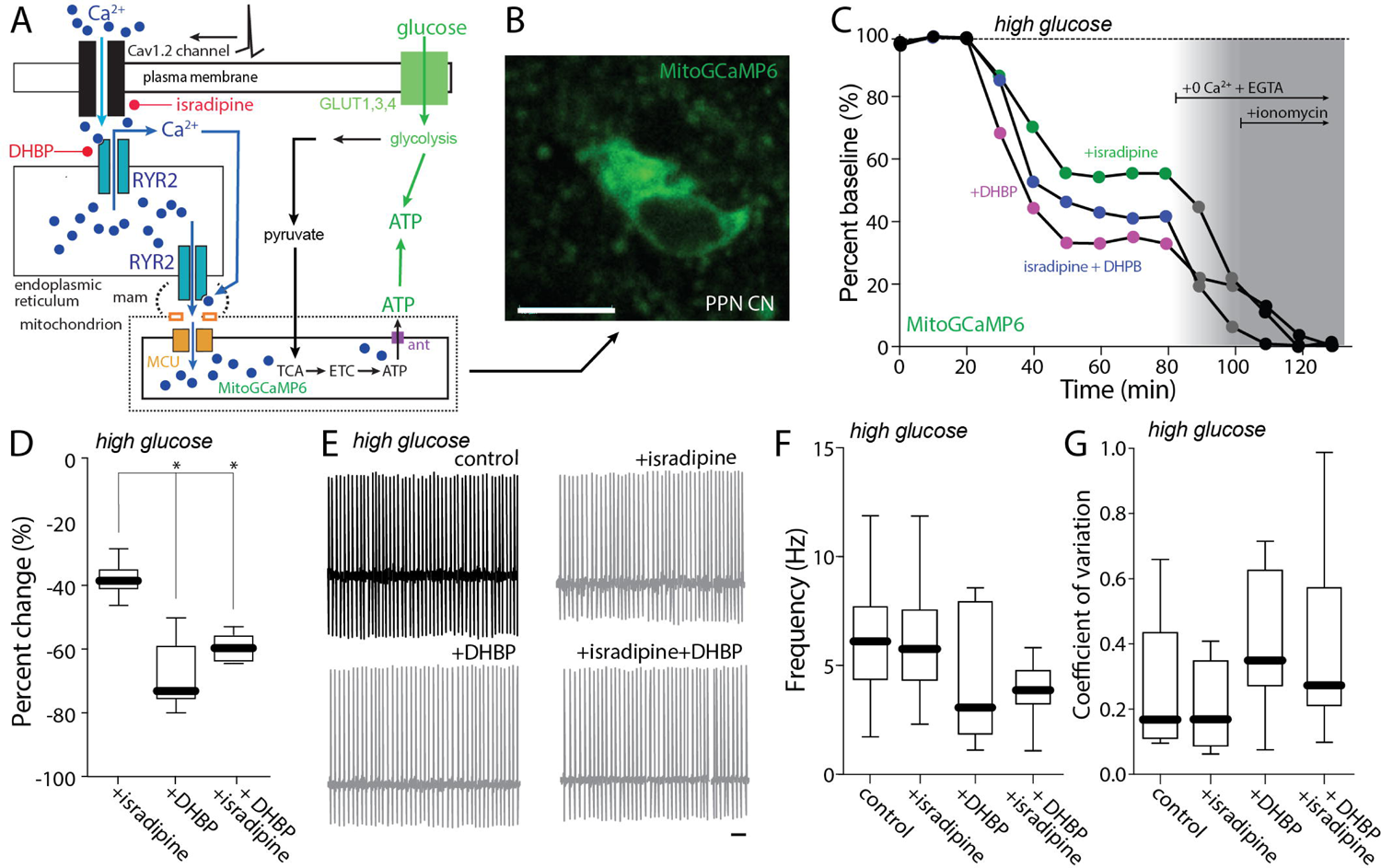
Cav1 block effect on pacemaking is mediated by control of mitochondrial matrix Ca2+ content. **A.** Cartoon illustrating the effect of Ca^2+^ entry through Cav1.2 channels on RYR Ca^2+^ release in a high-glucose (25 mM) aCSF. **B.** Representative micrograph of a recorded PPN CN expressing mitoGCaMP6. Bar: 10 μm. **C.** Representative examples of the percentage of mitochondrial Ca^2+^ in a PPN CN before and after addition of isradipine (500 nM), DHBP or both, 0 Ca^2+^ + EGTA and 0 Ca^2+^ + EGTA + ionomycin, all in a high-glucose (25 mM) aCSF. **D.** Percentage of change of mitochondrial Ca^2+^ from aCSF for isradipine, DHBP and both (Kruskal-Wallis, Interaction: *p<0.0001; Dunn’s multiple comparision, Isradipine-DHBP: *p=0.0016; Isradipine-Isradipine+DHBP: ^#^p=0.0520; n=6 in each group). **E.** Representative cell-attached recordings of PPN CNs without or with chronic isradipine (500 nM, 1 hr pre-incubation), chronic DHBP (100 μM, 1 hr pre-incubation), or both, in the presence of picrotoxin (100 mM) and CNQX (50 mM) in a high-glucose (25 mM) aCSF. Bar: 1 s. **F-G.** Firing frequency (**F**) and coefficient of variation (**G**) of PPN CNs with or without isradipine, DHBP, or both (**F:** Non-parametric one-way ANOVA, interaction NS; Dunn’s multiple comparision: control-Isradipine: NS; control-DHBP: NS; control-Isradipine+DHBP: NS; Isradipine-DHBP: NS; Isradipine-Isradipine+DHBP: NS; DHBP-Isradipine+DHBP: NS. **G:** Non-parametric one-way ANOVA, interaction NS; Dunn’s multiple comparision: control-Isradipine: NS; control-DHBP: NS; control-Isradipine+DHBP: NS; Isradipine-DHBP: NS; Isradipine-Isradipine+DHBP: NS; DHBP-Isradipine+DHBP: NS; n=9 in each group). Box plots indicate first and third quartiles, thick center lines represent medians, and whiskers indicate the range.

### Cav1 channels regulated mitochondrial ATP synthesis

Together, these data are consistent with the hypothesis that plasma membrane Cav1 Ca^2+^ channels are modulating pacemaking in PPN CNs by enhancing mitochondrial matrix Ca^2+^ content and the generation of ATP by OXPHOS. Indeed, Ca^2+^ entry into the mitochondrial matrix activates tricarboxylic acid cycle (TCA) enzymes and stimulates the production of reducing equivalents for the electron transport chain (ETC) and maintenance of the electrochemical gradient necessary for complex V conversion of ADP to ATP (Duchen, 1992; Kann et al., 2003; Yellen, 2018; Zampese & Surmeier, 2020). If this hypothesis is correct, cytosolic ATP concentration in pacemaking PPN CNs should be sensitive to disruption of the signaling pathway anchored by Cav1 channels. To test this prediction, a genetically encoded sensor for the ratio of ATP to adenosine diphosphate (ADP) – PercevalHR – was used (Tantama et al., 2013; Zampese et al., 2022). As described above, the PPN of ChAT-Cre/Ai14 mice were injected with an AAV carrying a Cre-dependent expression construct for PercevalHR, yielding cell-type specific expression in PPN CNs several week later (Fig. 5A-D). In ex vivo brain slices from these mice, Cav1 channels and RYRs were disrupted pharmacologically while monitoring cytosolic ATP/ADP ratio using 2PLSM (Fig. 5E). As predicted, inhibition of Cav1 Ca^2+^ channels or RYRs led to a roughly 40% drop in cytosolic ATP/ADP ratio (Fig. 5F); subsequent application of the complex V (ATP synthase) with oligomycin (10 µM) and then glycolysis with 2-deoxyglucose (3.5 mM) further decreased cytosolic ATP/ADP ratio. Using this dynamic range, the contribution of OXPHOS to the maintenance of cytosolic ATP/ADP ratio was estimated to be about 60% – a value very close to that of SNc dopaminergic neurons (Fig. 5G) (Zampese et al., 2022). On average, isradipine, DHBP and the combination produced 40% drop in the cytosolic ATP/ADP ratio, accounting for roughly two-thirds of the mitochondrial contribution (Fig. 5H).

**Figure 5:**
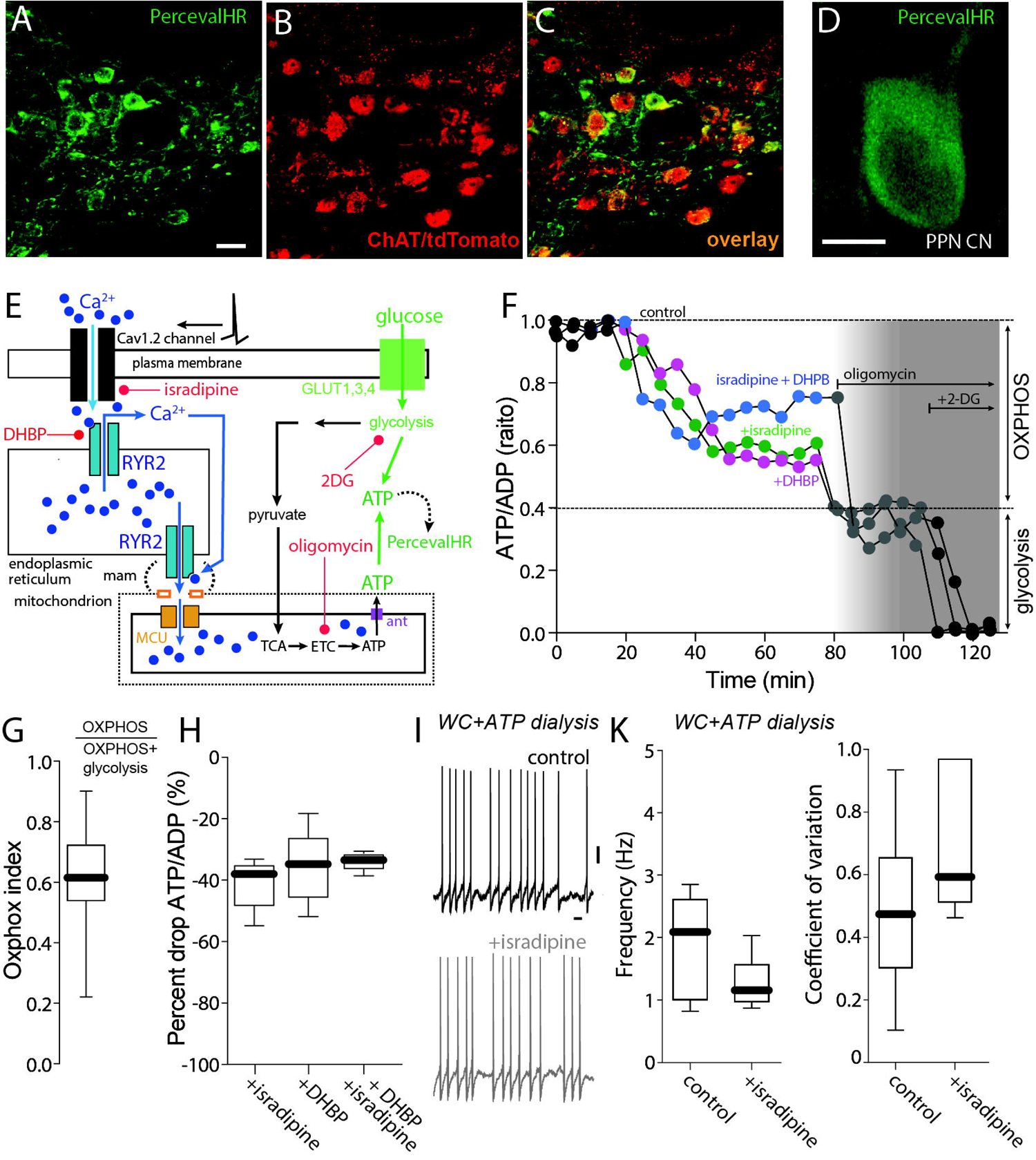
Cav1 channels regulate mitochondrial ATP synthesis. **A-C.** Representative micrograph from a ChAT-Cre/Ai14 mouse showing Perceval expression (green, **A**) in choline acetyltransferase immunolabeled neurons (ChAT, **B**) and the overlay of both markers (**C**). Bar: 20 μm. **D.** Representative micrograph of a recorded PPN CN expressing the PercevalHR probe. **E.** Cartoon illustrating the effect of Ca^2+^ entry through Cav1 channels and RYR Ca^2+^ release on mitochondrial ATP production and the action of isradipine and DHBP. **F.** Representative PercevalHR experiment estimating the effect of isradipine (500 nM) and DHBP (100 μM) on the contribution of mitochondria OXPHOS (inhibited by oligomycin) and glycolysis (inhibited by 2-DG) to the ATP/ADP ratio of the cell; the fluorescence measured for each of the two wavelengths (950 for ATP, 820 for ADP) is illustrated. **G.** Oxphox index of PPN CNs. **H.** Percentage of drop of ATP/ADP ratio from ACSF for isradipine, DHBP and both **I.** Representative whole-cell recordings of PPN CNs without or with chronic isradipine (500 nM, 1 hr pre-incubation). Bar: 1 s. **K.** Firing frequency and coefficient of variation of PPN CNs with or without isradipine (frequency: Mann-Whitney, NS; coefficient of variation: Mann-Whitney, NS; n=6 in each group). Box plots indicate first and third quartiles, thick center lines represent medians, and whiskers indicate the range.

Lastly, if the effect of isradipine on pacemaking was mediated by alterations in cytosolic ATP concentration, then nominally clamping ATP concentration with whole cell (WC) dialysis should blunt the effects. To test this prediction, visually identified PPN CNs were subjected to WC recording in ex vivo brain slices with a patch electrode containing 2 mM ATP. Spiking was slower and less regular in this configuration than in cell-attached recordings (cf. Fig. 2D,E, Fig. 3H-I and Fig. 5I,J). Nevertheless, in WC recordings, isradipine failed to have a significant effect on either rate or regularity (Fig. 5I,J).

### Activation of K-ATP K^+^ channels mediated the effect of isradipine on pacemaking

The experiments described to this point make the case that Cav1 Ca^2+^ channels participate in a feed-forward pathway that stimulates mitochondrial ATP production necessary for the maintenance of autonomous pacemaking. Cytosolic ATP concentration is important for a variety of processes that might impact pacemaking. One way in which a drop in cytosolic ATP concentration might slow pacemaking is by activation of plasma membrane K-ATP channels (Ashcroft, 2005). Interestingly, bath application of the K-ATP channel inhibitor glibenclamide (100 nM) had no effect on basal pacemaking in cell-attached recordings, but did significantly blunt the effect of isradipine on spiking rate and regularity (Fig. 6A-C). In RNASeq and qPCR profiling of RiboTag isolated mRNA from PPN CNs, both Kir6.1/2 and SUR1 transcripts, which code for K-ATP channel subunits, were detected (Fig. 6D). Together, these data support a feed-forward model of metabolic control (Fig. 6E).

**Figure 6:**
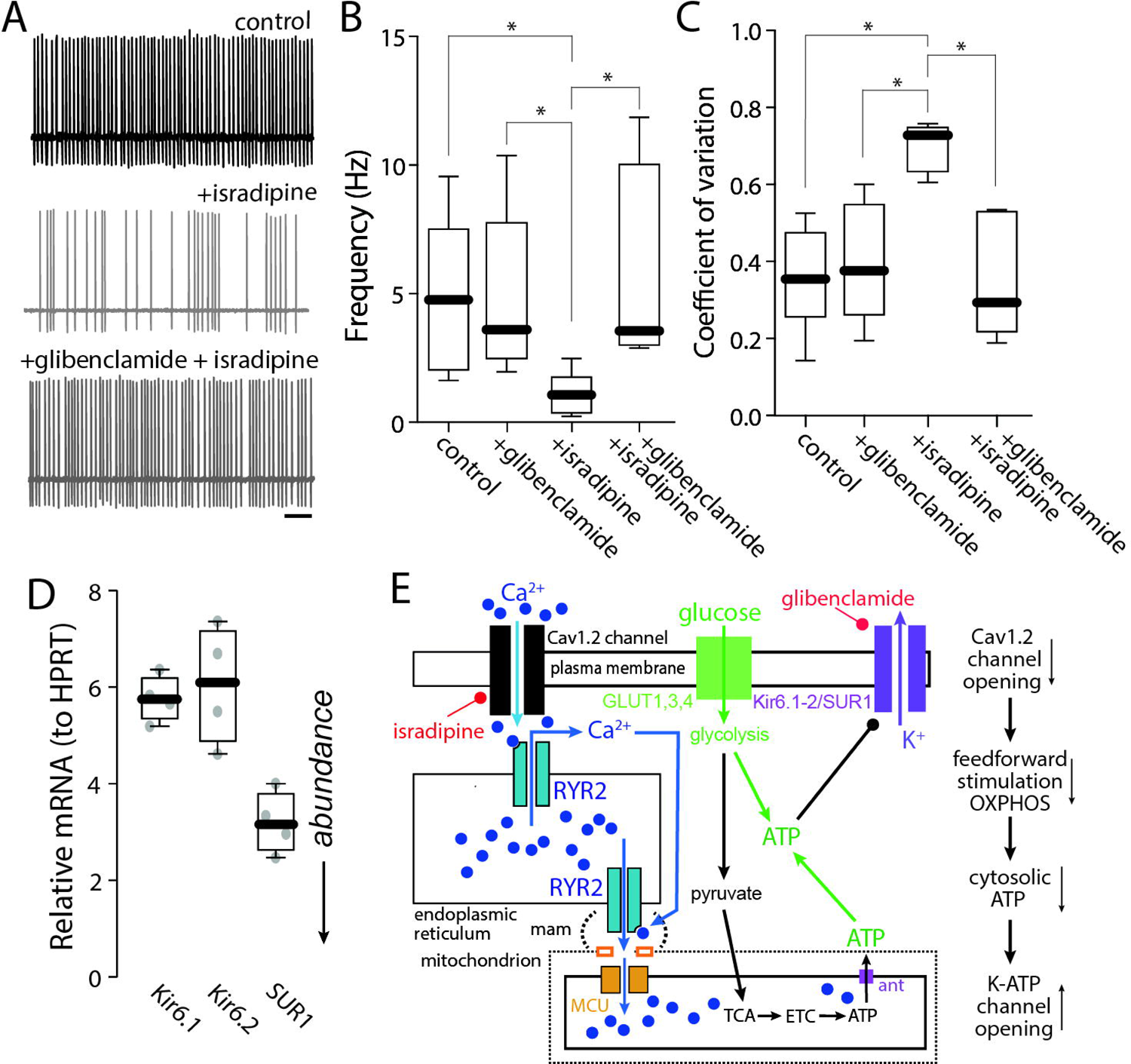
Activation of K-ATP K^+^ channels mediated the effect of isradipine on pacemaking. **A.** Representative cell-attached recordings of PPN CNs without or with chronic isradipine (500 nM, 1 hr pre-incubation), before and after application of Glibenclamide (100 nM) in the presence of the synaptic blockers picrotoxin and CNQX. Bar: 1 s. **B-C.** Firing frequency (**B**) and coefficient of variation (**C**) of PPN CNs with or without isradipine, before and after application of Glibenclamide (**B:** Non-parametric one-way ANOVA, interaction *p=0.0064; Dunn’s multiple comparision: control - +Glibenclamide: NS; control - Isradipine: *p<0.05; control - Isradipine+Glibenclamide: NS; Isradipine - +Glibenclamide: *p<0.05; Isradipine+Glibenclamide - +Glibenclamide: NS; Isradipine - Isradipine+Glibenclamide: *p<0.05. **C:** Non-parametric one-way ANOVA, interaction *p=0.0087; Dunn’s multiple comparision: control - +Glibenclamide: NS; control - Isradipine: *p<0.05; control - Isradipine+Glibenclamide: NS; Isradipine - +Glibenclamide: *p<0.05; Isradipine+Glibenclamide - +Glibenclamide: NS; Isradipine - Isradipine+Glibenclamide: *p<0.05; n=7 in each group). Box plots indicate first and third quartiles, thick center lines represent medians, and whiskers indicate the range. **D.** Quantification of RNA abundance of Kir6.1, Kir6.2 ans SUR subunits. **E.** Cartoon illustrating the effect of Ca^2+^ entry through Cav1 channels and RYR Ca^2+^ release on K_ir6_/SUR1 channels.

### Feed-forward stimulation of mitochondria increased oxidant stress

One of the downsides of feed-forward stimulation of mitochondrial OXPHOS is that it leads to increased production of reactive oxygen species (ROS). ROS damage lipids, proteins and DNA, potentially causing mitochondrial damage and increase autophagic load, as well as aging related decline in mitochondrial function and cellular vitality (Lionaki et al., 2015). To determine if mitochondrial oxidant stress was elevated in PPN CNs by engagement of feed-forward stimulation of OXPHOS, a genetically encoded, mitochondrially targeted redox sensor (mito-roGFP) was used in conjunction with 2PLSM in ex vivo brain slices. To limit mito-roGFP expression to PPN CNs, the PPN of ChAT-Cre/Ai14 mice were injected with an AAV carrying a Cre-dependent mito-roGFP expression construct. Several weeks after injection, this yielded robust mito-roGFP expression in PPN CNs (Fig. 7A-D). Using a well-established calibration strategy (Guzman et al., 2010; Zampese et al., 2022), the basal oxidation of mito-roGFP was estimated to be around 35% (Fig. 7E,F), somewhat lower than that seen in SNc dopaminergic neurons (Guzman et al., 2010; Zampese et al., 2022). Bath application of isradipine significantly lowered the level of mitochondrial oxidation, consistent with the hypothesis that feed-forward stimulation of OXPHOS increased oxidant stress levels in PPN CNs (Fig. 7E,F). A graphical summary of these results is presented in Figure 7G.

**Figure 7:**
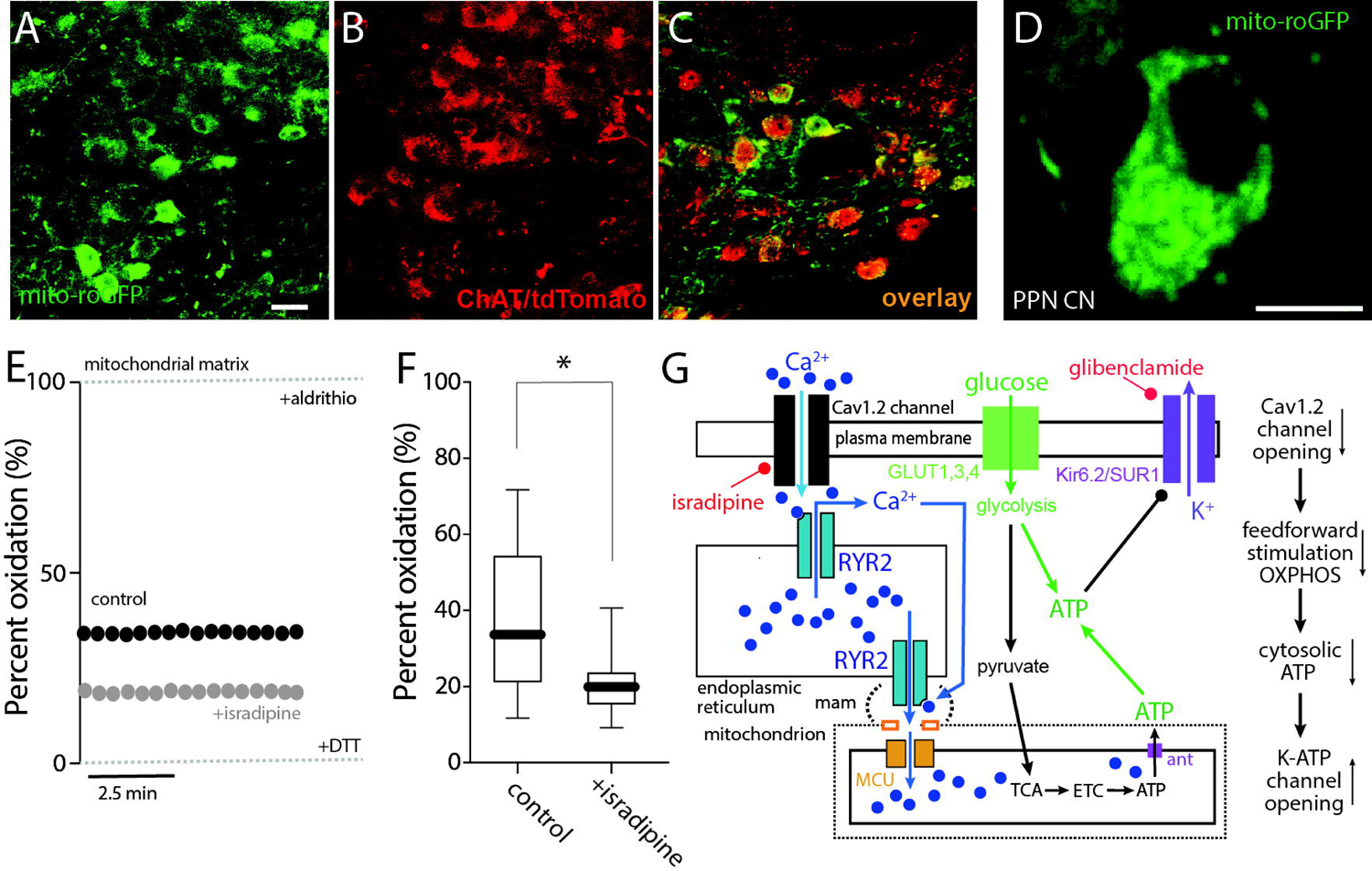
Feed-forward stimulation of mitochondria increased oxidant stress. **A-C.** Representative micrograph from a ChAT-Cre/Ai14 mouse showing mito-roGFP expression (green, **A**) in choline acetyltransferase immunolabeled neurons (ChAT, **B**) and the overlay of both markers (**C**). Bar: 20 μm. **D.** Representative micrograph of a recorded PPN CN expressing mito-roGFP. Bar: 10 μm. **E.** Representative examples of the percentage of oxidation of a PPN CN in control aCSF or with isradipine, expressed as relative oxidation compared to the fully reduced and fully oxidized states obtained upon application of dithiothreitol (DTT; 2 mM) and aldrithiol (Ald; 200 μM), indicated by dashed lines. **F.** Percentage of oxidation of PPN CNs in the absence or presence of isradipine (Mann-Whitney, *p=0.0234; n=12-14 in each group). Box plots indicate first and third quartiles, thick center lines represent medians, and whiskers indicate the range. **G.** Cartoon summarizing the effect of Ca^2+^ entry through Cav1 channels and RYR Ca^2+^ release on ROS generation.

## Discussion

Our studies demonstrate that PPN CNs rely upon feed-forward stimulation of mitochondrial OXPHOS to provide the ATP necessary to sustain regular, autonomous pacemaking. The feed-forward Ca^2+^ signaling pathway relied upon plasma membrane L-type Ca^2+^ channels with a Ca_v_1.2 pore-forming subunit, ER RYRs and Ca^2+^ entry into mitochondria – a well-known mode of OXPHOS stimulation. Disrupting this spike-dependent signaling led to a drop in cytosolic ATP/ADP ratio and activation of K-ATP channels that slowed pacemaking rate and decreased its regularity. As in other neurons at-risk in PD, engagement of this feed-forward signaling pathway increased mitochondrial oxidant stress, providing a mechanistic linkage to aging-related loss of mitochondrial function and aggregation of aSYN, both of which are PD risk factors.

### PPN CNs rely upon Cav1.2 Ca^2+^ channels for feed-forward bioenergetic signaling

All cells must match their energy needs with their production of ATP. In neurons, ATP is generated from glucose by glycolysis and by mitochondrial metabolism of glycolytic by-products – pyruvate and malate. Mitochondria use reducing equivalents generated by pyruvate and malate metabolism to drive the electron transport chain (ETC), which creates the electrochemical gradient used by mitochondrial complex V (MCV or ATP synthase) to convert ADP to ATP. Given that mitochondrial OXPHOS generates an order of magnitude more ATP per glucose molecule than does glycolysis, they are critical to bioenergetically demanding and substrate limited neurons (Duchen, 1992; Kann et al., 2003; Pfeiffer et al., 2001; Yellen, 2018; Zampese & Surmeier, 2020). Mitochondrial ATP generation from glycolytic metabolites is regulated by both feedback and feed-forward control. MCV is subject to substrate inhibition so that as cytosolic ATP/ADP ratio falls, MCV is dis-inhibited (Zampese et al., 2022). Complementing this feedback control mechanism is a feed-forward signaling pathway that relies upon Ca^2+^(Griffiths & Rutter, 2009). The availability of reducing equivalents for the ETC is regulated by Ca^2+^ reaching the mitochondrial intermembrance space and matrix (Gellerich et al., 2013; Szibor et al., 2020): intermembrane space Ca^2+^ stimulates the malate-aspartate shuttle, whereas matrix Ca^2+^ stimulates pyruvate metabolism by the tricarboxylic acid (TCA) cycle. In muscle, the Ca^2+^ driving mitochondrial metabolism is derived from opening of plasma membrane L-type Ca^2+^ channels that trigger RYR-dependent Ca^2+^ release from intracellular stores (Díaz-Vegas et al., 2018; Viola et al., 2009). Each element in this cascade – L-type Ca^2+^ channels, RYRs and mitochondria – are positioned physically close to one another to maximize the efficiency of the couple (Díaz-Vegas et al., 2018; Viola et al., 2009). Because spikes are excellent predictors of global bioenergetic demand in neurons, making feed-forward control dependent upon L-type Ca^2+^ channels that are efficiently opened by spikes and manifest relatively little voltage-dependent inactivation makes obvious biological sense (Duchen, 1992; Kann et al., 2003; Yellen, 2018). Indeed, in SNc dopaminergic neurons where bioenergetic control mechanisms have been studied in-depth, this feed-forward control mechanism is clearly engaged and important to matching bioenergetic demand to ATP supply (Zampese et al., 2022).

In PPN CNs, all of the feed-forward signaling elements seen in SNc dopaminergic neurons are in place. In ex vivo brain slices, where PPN CNs were autonomously pacemaking, mitochondrial matrix Ca^2+^ content (measured using a matrix targeted variant of GCaMP6) was lowered by pharmacological inhibition of L-type Ca^2+^ channels or RYRs. Importantly, the lowering of matrix Ca^2+^ content achieved by inhibition of L-type channels or RYRs was not additive, but occlusive, arguing that they were part of the same signaling cascade. Consistent with this interpretation, cytosolic ATP/ADP ratio (measured using cytosolic PercevalHR) was lowered by inhibiting L-type Ca^2+^ channels or by inhibiting RYRs and the effect of combined inhibition was similar to that of inhibiting either one alone.

One of the apparent differences between PPN CNs and SNc dopaminergic neurons was the subtype of L-type Ca^2+^ channel involved. Profiling of PPN CNs mRNA harvested using the RiboTag method revealed robust expression of CACNA1C, but not CACNA1D – arguing that L-type channels with a Cav1.2 pore-forming subunit were the principal contributors to feed-forward control. In SNc dopaminergic neurons, Cav1.3 channels play a predominant role. These two channel types differ primarily in their voltage-dependence with Cav1.2 channels being activated by membrane depolarization above spike threshold, whereas Cav1.3 channels are activated at sub-threshold potentials. The reasons for this difference are not clear. Given that PPN CNs robustly expressed mRNA coding for low threshold Cav1.3 Ca^2+^ channels argues that the difference was not due to a physiological design that broadly limited Ca^2+^ entry at sub-threshold membrane potentials. Given the privileged linkage of L-type channels to bioenergetic control, it could be the energetic demands on SNc dopaminergic neurons are greater than those of PPN CNs, requiring stimulation of mitochondria for more of the pacemaking duty cycle.

That said, all of the factors governing a neuron’s bioenergetic ledger sheet have not been rigorously characterized. Much has been made of the bioenergetic needs associated with regenerative spiking, which undoubtedly is important to neurons (Duchen, 1992; Kann et al., 2003; Yellen, 2018). However, a neuron’s somatodendritic region has a variety of other demands. For example, many of the anabolic and catabolic processes associated with proteostasis, like those associated with maintaining an axonal arbor with thousands of transmitter release sites, reside in the somatodendritic region (Lottes & Cox, 2020; Pacelli et al., 2015). Although the axonal arbor of PPN CNs is unusually large (Mena-Segovia et al., 2008), whether it poses as much of a burden as that of a ventral tier SNc dopaminergic neuron remains to be determined. Another peculiar feature of SNc dopaminergic neurons that may energetically stress the somatodendritic region is the local release of dopamine; PPN CNs are not known to release ACh from their dendrites.

### Basal pacemaking required feed-forward stimulation of OXPHOS

A key inference to be drawn from our studies is that PPN CNs require feed-forward stimulation of mitochondrial OXPHOS to sustain basal pacemaking. When PPN CNs were bathed in physiological concentrations of glucose (3.5 mM), inhibition of L-type Ca^2+^ channels led to a drop in cytosolic ATP/ADP ratio and engagement of K-ATP K^+^ channels that lowered pacemaking rate and regularity. Elevating extracellular glucose to increase glycolytic ATP production or dialysis with an ATP-containing internal solution blunted the effects of inhibiting L-type channels, arguing that the effects on spike generation were bioenergetic, not electrogenic. On the face of it, this dependence seems at odds with the situation in SNc dopaminergic neurons where inhibition of L-type channels had little or no effect on basal pacemaking recorded in cell-attached or perforated patch mode (Roeper, 2013; Zampese et al., 2022). However, this difference does not reflect any significant variation in the feed-forward control of mitochondria or the bioenergetic demands of pacemaking. Rather, it simply reflects the higher basal pacemaking rate in PPN CNs, which is roughly twice that of SNc dopaminergic neurons recorded in the same way, at the same temperature. Increasing the basal pacemaking rate of SNc dopaminergic neurons into the 4-5 Hz range typical of PPN CNs revealed a clear dependence upon L-type Ca^2+^ channel stimulation of mitochondrial OXPHOS (Zampese et al., 2022).

Why PPN CNs have a higher basal pacemaking rate that SNc dopaminergic neurons is unclear. This rate is likely to reflect a combination of factors. As mentioned above, one is likely to be the bioenergetic demands linked to somatodendritic proteostasis. Another is likely to be the network requirements for basal ACh release in regions innervated by PPN CNs.

### Translational implications for PD

The loss of PPN CNs is associated with both motor and non-motor symptoms of PD (Chambers et al., 2019; Muslimović et al., 2008; Sethi, 2008; Zweig et al., 1987). Our results provide insight into their selective vulnerability. The reliance of PPN CNs on feed-forward stimulation of mitochondrial OXPHOS to meet basal bioenergetic needs comes at the cost of sustained elevation in oxidant stress. This is a common feature of other at-risk neurons, including SNc dopaminergic neurons, locus coeruleus (LC) adrenergic neurons and dorsal motor nucleus of the vagus (DMV) cholinergic neurons (Goldberg et al., 2012; Guzman et al., 2010; Sanchez-Padilla et al., 2014; Zampese et al., 2022). Sustained oxidant stress is associated with a variety of potential drivers of PD pathogenesis, including loss of mitochondrial function and aggregation of alpha-synuclein (aSYN). Indeed, post-mortem analysis of PD brains shows that PPN CNs manifest Lewy pathology (LP) rich in misfolded aSYN (Hirsch et al., 1987; Jellinger, 1988).

Although it remains uncertain whether PPN LP can arise in a cell autonomous manner, there are compelling reasons to think that misfolded forms of aSYN can propagate through neural networks (Henderson et al., 2019). In healthy mouse models, local deposition of misfolded, pre-formed fibrillar (PFF) forms of aSYN (mimicking propagation) leads to the formation of Lewy-like intracellular inclusions specifically in PPN CNs, not in neighboring glutamatergic or GABAergic neurons – both of which are relatively resistant in PD (Henrich et al., 2020). As activity-dependent macropinocytosis contributes to neuronal uptake of extracellular aSYN PFFs (Okuzumi et al., 2018), autonomous pacemaking of PPN CNs may contribute to this selectivity. Basal mitochondrial oxidant stress might promote the misfolding of endogenous aSYN and the growth of cytoplasmic LP; it might also and increase the catabolic burden on PPN CNs by damaging other proteins, lipids and deoxyribonucleic acid (DNA) (Lionaki et al., 2015). In SNc dopaminergic neurons, which have a similar basal mitochondrial oxidant stress level, mitophagy is elevated compared to neighboring ventral tegmental area dopaminergic neurons that are relatively resistant in PD (Guzman et al., 2010). If this feature generalizes to PPN CNs, it may reduce spare proteostatic capacity and limit their ability to deal with exogenous stressors, like misfolded forms of aSYN.

The reliance of PPN CNs on L-type Ca^2+^ channels to drive mitochondrial respiration and oxidant stress also has translational implications. As shown here, reducing the gain of the feed-forward pathway linking spiking to mitochondrial OXPHOS by partial inhibition of L-type channels with dihydropyridines decreases oxidant stress. This is true not just in PPN CNs but in other neurons known to be selectively vulnerable in PD (Surmeier, Obeso, et al., 2017). Forcing neurons to rely more on glycolysis and less on mitochondrial OXPHOS (González-Rodríguez et al., 2021) should increase their resilience and slow degeneration. Indeed, the early, systemic use of dihydropyridines is strongly associated with a dose-dependent reduction in the risk of developing PD (Gudala et al., 2015; Lang et al., 2015). Moreover, in early stage PD patients, oral administration of a sustained release format of isradipine appears to slow disease progression (Surmeier, Schumacker, et al., 2017). Although a Phase 3 clinical trial with an immediate release format of isradipine failed to reach its clinical goals, the lack of sustained inhibition of Cav1 channels with this drug format makes the interpretation of this result problematic, particularly given evidence of disease slowing in that subset of patients with slower systemic drug clearance (Venuto et al., 2021).

Given that dihydropyridines have a higher affinity for Cav1.2 Ca^2+^ channels than those with a Cav1.3 pore (Liss & Striessnig, 2019; Ortner et al., 2017), they should be more effective in slowing the loss of PPN CNs than SNc dopaminergic neurons. It would be of considerable interest to know whether individuals taking dihydropyridines that do manifest parkinsonian motor symptoms have fewer PPN-linked gait, balance and sleep deficits than those that are unmedicated. The reliance of PPN CNs on Cav1.2 channels also would limit the therapeutic impact of Cav1.3 channel selective inhibitors in development (Cooper et al., 2021).

## Methods

### Animals

Mice allowing the identification of PPN CNs through cell type-specific expression of the fluorescent protein tdTomato (ChAT-Cre/Ai14, for simplicity) were obtained by crossing ChAT-Cre (B6;129S6-Chattm2(cre)Lowl/J, Stock 6410,The Jackson Laboratories (Rossi et al., 2011) and Rosa-CAG-LSL-tdTomato-WPRE mice (B6.CgGt(ROSA)26Sortm14(CAG-tdTomato)Hze/J, stock 7914,The Jackson Laboratories (Madisen et al., 2010). 2-month-old female and male mice were used. All mice were bred in-house and used with the approval by Northwestern University Animal Care and Use Committee and in accordance with the National Institutes of Health (NIH) Guide for the Care and Use of Laboratory Animals. Mice were group-housed with food and water *ad libitum* on a 12-hour light/dark cycle with temperatures of 65° to 75°F and 40 to 60% humidity.

### Virus generation

PCR-amplified sequences for GCaMP6s (Chen et al., 2013), 2mt-GCaMP6m (Logan et al., 2014), PercevalHR (Tantama et al., 2013), and mito-roGFP (Dooley et al., 2004) were subcloned into Eco RI and Sal I restriction sites of the pFB-TH-SV40 vector and packaged into recombinant AAVs, serotype 9, titers ∼2×1013 vp (viral particles)/ml (Virovek).

### Stereotaxic surgery

An isoflurane precision vaporizer (Smiths Medical PM) was used to anesthetize mice. Mice were then placed on a stereotaxic frame (David Kopf Instruments), with a Cunningham adaptor (Harvard Apparatus) to maintain anesthesia delivery during surgery. The skull was exposed, and a small hole was drilled at the desired injection site. The following stereotaxic coordinates were used to inject into PPN: 4.4 and 1.25 mm posterior and lateral to bregma, at a depth of 3.5 mm from dura. The Allen Mouse Brain Atlas, online version 1, 2008 (https://atlas.brain-map.org/) was used as a reference for the coordinates and generating diagrams. For each mouse, the distance between bregma and lambda was calculated and used to adjust the coordinates. For AAV injections, ∼200 nl of viral vector was delivered using a glass micropipette (Drummond Scientific) pulled with a P-97 glass puller (Sutter Instruments). All surgeries were performed bilaterally. Experiments were performed after at least 15 postoperative days; tissue collection for RiboTag was performed 4 weeks after injection.

### Ex vivo slice preparation

Mice were anesthetized with a mixture of ketamine (50 mg/kg) and xylazine (4.5 mg/kg) and transcardially perfused with ice-cold modified artificial cerebrospinal fluid (aCSF) (“slicing solution,” containing 49.14 mM NaCl, 2.5 mM KCl, 1.43 mM NaH_2_PO_4_, 25 mM NaHCO_3_, 25 mM glucose, 99.32 mM sucrose, 10 mM MgCl_2_, and 0.5 mM CaCl_2_); after decapitation, the brain was removed and sectioned in 250 um-thick parasagittal slices with a 20° angle, placing the brain in a 20° wedge of agarose gel, using a vibratome (VT1200S, Leica Microsystems) in the same slicing solution. Slices were transferred into a holding chamber containing recording aCSF (“physiological glucose aCSF,” containing 135.75 mM NaCl, 2.5 mM KCl, 1.25 mM NaH_2_PO_4_, 25 mM NaHCO_3_, 2 mM CaCl_2_, 1 mM MgCl_2_, and 3.5 mM glucose). Slices were allowed to recover for 30 min at 34°C and then kept at room temperature for at least 15 min before starting the experiments. For specific experiments, aCSF was modified by replacing glucose (3.5 mM) with the same concentration of 2-DG (3.5 mM, Sigma-Aldrich). High-glucose aCSF contained 125 mM NaCl, 2.5 mM KCl, 1.25 mM NaH_2_PO_4_, 25 mM NaHCO_3_, 2 mM CaCl_2_, 1 mM MgCl_2_, and 25 mM glucose. All solutions were pH 7.4 and ∼310 mOsm, and continually bubbled with 95% O_2_ and 5% CO_2_.

For isradipine preincubation experiments, after recovery, slices were transferred to a holding chamber containing physiological glucose aCSF with 500 nM isradipine for at least 1 hour before starting the experiments. All *ex vivo* brain slice experiments were conducted at 34° C.

### 2PLSM optical workstation

The foundation of the system is the Olympus BX-51WIF upright microscope with an LUMPFL 60×/1.0 NA (numerical aperture) water-dipping objective lens and an Ultima dual excitation-channel scan head (Bruker Nano Fluorescence Microscopy Unit). The automation of the XY stage motion, lens focus, and manipulator XYZ movement was provided by FM-380 shifting stage, axial focus module for Olympus scopes, and manipulators (Luigs & Neumann). Cell visualization and patching were made possible by a variable magnification changer, calibrated to 2× (100-m field of view) as defined by the LSM bright-field transmission image, supporting a 1-megapixel USB3.0 complementary metal-oxide semiconductor (CMOS) camera (DCC3240M, Thor Labs) with ∼30% quantum efficiency around 770 nm. Olympus NIR-1 band-pass filter, 770 nm/100 nm, and Manager (Edelstein et al., 2010) software were used with the patch camera. The electrical signals were sent and collected with a 700B patch clamp amplifier, and MultiClamp Commander software with computer input and output signals was controlled by Prairie View 5.3-5.5 using a National Instruments PCI-6713 output card and PCI-6052e input card. The 2P excitation (2PE) imaging source was a Chameleon Ultra1 series tunable wavelength (690 to 1040 mm, 80 MHz, ∼250 fs at sample) Ti: sapphire laser system (Coherent Laser Group); the excitation wavelength was selected on the basis of the probe being imaged (see below). Each imaging laser output is shared (equal power to both sides) between two optical workstations on a single anti-vibration table (TMC). Workstation laser power attenuation was achieved with two Pockels’ cell electro-optic modulators (models M350-80-02-BK and M350-50-02-BK, Con Optics) controlled by Prairie View 5.3-5.5 software. The two modulators were aligned in series to provide enhanced modulation range for fine control of the excitation dose (0.1% steps over 5 decades), to limit the sample maximum power, and to serve as a rapid shutter during line scan or time series acquisitions. The 2PE generated fluorescence emission was collected by non–de-scanned photomultiplier tubes (PMTs). Green channel (490 to 560 nm) signals were detected by a Hamamatsu H7422P-40 select GaAsP PMT. Red channel (580 to 630 nm) signals were detected by a Hamamatsu R3982 side on PMT. Dodt-tube–based transmission detector with Hamamatsu R3982 side on PMT (Bruker Nano Fluorescence) allowed cell visualization during laser scanning. Scanning signals were sent and received by the National Instruments PCI-6110 analog-to-digital converter card in the system computer (Bruker Nano Fluorescence). All XY images were collected with a pixel size of 0.195 m, a pixel dwell time of 12 s, and a frame rate of 3 to 4 fps (frames per second).

### Ex vivo Ca^2+^ 2PLSM measurements

Slices were transferred to a recording chamber and continuously perfused with aCSF at 32° to 33°C. A laser wavelength of 920 nm was used for mito-GCaMP6 experiments. Alternatively, brightness over time (BOT) continuous measurements were collected during experiments with acute stimulation. To calibrate the dynamic range of mito-GCaMP6 probe in the experiments, slices were perfused with Ca2+-free aCSF (CaCl2 was substituted with MgCl_2_, for a total of 3 mM MgCl_2_) + 0.5 mM EGTA and 1 uM ionomycin to obtain Fmin, followed by aCSF (with total 3 mM CaCl2) + 1 uM ionomycin to obtain Fmax. Time series and BOT data were analyzed offline, and fluorescence measurements in multiple regions of interest (ROIs) were evaluated, with a separate background ROI value subtracted. Measurements are presented as F - Fmin, where Fmin is the minimum fluorescence obtained in Ca^2+^-free aCSF + ionomycin. Semiquantitative estimations of Ca^2+^ levels were calculated as the percentage of the dynamic range of the probe (% of MAX).

### Ex vivo redox 2PLSM measurements

Slices were transferred to a recording chamber and continuously perfused with aCSF at 32° to 33°C. Mito-roGFP experiments were performed using an excitation wavelength of 920 nm, as previously described (Guzman et al., 2010). Briefly, time series images were acquired with 60 frames obtained over ∼20 s. The dynamic range of the probe was determined by bath applying 2 mM dithiothreitol, a reducing agent, followed by 200 uM aldrithiol, an oxidizing agent, to determine the fluorescence intensities of the probe at minimal and maximal oxidation for each cell. Time series were analyzed offline, multiple ROIs were measured, the background ROI value was subtracted, and then the relative oxidation was calculated.

### Ex vivo ATP/ADP 2PLSM measurements

Slices were transferred to a recording chamber and continuously perfused with aCSF at 32° to 33°C. For the excitation ratio probe PercevalHR, ATP/ADP ratio measurements of two excitation wavelengths, 950 nm for ATP and 820 nm for ADP, were used in rapid succession for each acquisition time point and two time series of five frames (1.25 to 1.7 s long) were acquired for each wavelength. Time series were analyzed offline, a cytosolic and a background ROIs were measured, the background signal was subtracted, and the ATP/ADP ratio (950:820) was calculated for each ratio pair time point. The contribution of mitochondria to the bioenergetic status of each cell (OXPHOS index) was estimated comparing the drop in the PercevalHR ATP/ADP ratio induced by bath application of 10 uM oligomycin (Sigma-Aldrich) and the minimum ratio obtained in a modified aCSF where glucose was substituted with the nonhydrolyzable 2-DG plus 10 uM oligomycin. For experiments with other pharmacological manipulations, the percentage of drop in ATP/ADP ratio is presented as R - Rmin, where R is the ratio measured at the time point of interest and Rmin is the minimum ratio obtained in 2-DG aCSF + 10 uM oligomycin, normalized to the control situation.

### Electrophysiology and Fura-2 2PLSM Ca^2+^ imaging

Slices were transferred to a recording chamber on an Olympus BX51 upright microscope and perfused with oxygenated aCSF as described above. Patch pipettes were pulled from thick-walled borosilicate glass on a Sutter P-1000 puller; pipette resistance was 2.5 to 5 megaohms.

Electrophysiological recordings were obtained using a MultiClamp 700B amplifier. For most recordings, electrophysiological signals were filtered at 1 to 4 kHz and digitized at 10 kHz. The liquid junction potential was not compensated. For experiments performed in whole-cell configuration, pipettes were filled using a potassium-based internal solution containing 126 mM KMeSO4, 14 mM KCl, 10 mM Hepes, 1 mM EGTA, 0.5 mM CaCl_2_, and 3 mM MgCl_2_; pH was adjusted to 7.25 to 7.3 with KOH, osmolarity 280 to 300 mOsm. For cell attached configuration, pipettes were filled with aCSF.

For Fura-2 imaging, conventional tight-seal whole-cell patchclamp recordings were made from PPN cholinergic neurons. The patch electrode solution contained 135 mM KMeSO_4_, 5 mM KCl, 5 mM Hepes, 10 mM Na_2_PCr (phosphocreatine disodium), 2 mM ATP-Mg, and 0.5 mM guanosine triphosphate (GTP)–Na, pH ∼ 7.3 and osmolarity 290 to 300 mOsm, with addition of Fura-2 (100 uM) and Alexa Fluor 594 hydrazide (25 uM; Thermo Fisher Scientific). A 780-nm laser wavelength was used to perform 2PLSM Fura-2 Ca^2+^ imaging using line scan acquisitions with 0.195-m pixels and 12-s dwell time, and [Ca^2+^] was estimated using the following equation: [Ca^2+^] = Kd [1 - f(t)/fmax]/[f(t)/fmax - 1/Rf], as previously described (25). A Kd of 200 nM and Rf of 20 were assumed and used.

### Pharmacology

Drugs were prepared as stock solutions, diluted in aCSF immediately before use and applied through the perfusion system. The following stock solvents and final concentrations were used: distilled H20 for DHBP (Tocris, 100 uM), dithiothreitol (Invitrogen, 2 mM) and aldrithiol (Invitrogen, 200 uM); DMSO for CNQX (Tocris, 50 uM), picrotoxin (Sigma-Aldrich, 100 uM), Isradipine (Sigma-Aldrich, 500 nM), Glibenclamide (Tocris, 100 nM), ionomycin (Tocris, 1 uM) and oligomycin A (Sigma-Aldrich, 10 uM).

### Immunofluorescence and confocal imaging

For confirmation of ChAT expression in tdTomato positive PPN neurons and viral vectors localization in PPN, mice with Ribotag-GFP, Mito-GCaMP6, Perceval and mito-roGFP vectors were transcardially perfused for brain fixation. Fixed tissue was prepared by perfusing terminally anesthetized mice (ketamine 50 mg/kg and xylazine 4.5 mg/kg) with phosphate-buffered saline (PBS; Sigma-Aldrich) immediately followed by 4% paraformaldehyde (PFA; diluted in PBS from a 16% stock solution; Electron Microscopy Sciences). The brain was then removed and transferred into PFA solution for 4 hours before being thoroughly rinsed and stored in PBS at 4°C. Fixed brains were then sectioned into 80-um-thick parasagittal slices with a 20° angle, placing the brain in a 20° wedge of agarose gel on a Leica VT1200S vibratome and collected in PBS.

For ChAT immunofluorescence, PPN containing sections were bathed in a permeabilization solution (0.5% Triton X-100 in PBS) for 15 min, quickly washed with PBS, and then bathed in blocking solution [10% normal goat serum (NGS) and 0.25% Triton X-100 in PBS] for 1 hour. Anti-ChAT primary antibody (goat anti-ChAT; millipore AB 144), diluted 1:1000 in the same blocking solution, was applied overnight at 4°C. Subsequently, sections were washed three times in PBS and then incubated in the secondary antibody with green fluorophores (Alexa Green anti-goat, Life Technologies) diluted (1:500) in PBS for 1 hour at room temperature. After three washes in PBS, sections were prepared for image acquisition. In any case, PPN containing slices were transferred on microscopy slides (VWR) and allowed to dry and mounted with ProLong Diamond (Thermo Fisher Scientific) and #1.5 glass coverslips (VWR). Mounted sections were stored at 4°C until imaged with an Olympus FV10i confocal laser scanning microscope, using 10×/0.4 (air) or 60×/1.35 (oil) objective. FIJI (NIH) was used to adjust images for brightness, contrast, and pseudo-coloring.

### RiboTag profiling

AAVs for expression of RiboTag under a cre-dependent promoter (AAV9-EF1a-DIO-Rpl22l1-Myc-DDK-2A-tdTomato-WPRE, titers 2.24 × 1013 viral genomes/ml) were stereotaxically injected into the PPN. Four weeks after injection, mice were sacrificed and the PPN tissue expressing RiboTag was dissected and frozen at −80°C. RiboTag immunoprecipitation was carried out as previously described (Heiman M et al. Cell type-specific mRNA *purification by translating ribosome affinity purification (TRAP) Nat Protoc. 9, 1282-1291, 2014*). Briefly, tissue was homogenized in cold homogenization buffer [50 mM tris (pH 7.4), 100 mM KCl, 10 mM MgCl2, 1 mM dithiothreitol, cyclo-heximide (100 mg/ml), protease inhibitors, recombinant ribonuclease (RNase) inhibitors, and 1% NP-40]. Homogenates were centrifuged at 10,000*g* for 10 min, and the supernatant was collected and precleared with protein G magnetic beads (Thermo Fisher Scientific) for 1 hour at 4°C, under constant rotation. Immunoprecipitations were carried out with anti-Flag magnetic beads (Sigma-Aldrich) at 4°C overnight with constant rotation, followed by four washes in high-salt buffer [50 mM tris (pH 7.4), 350 mM KCl, 10 mM MgCl2, 1% NP-40, 1 mM dithiothreitol, and cycloheximide (100 mg/ml)]. RNA was extracted using RNeasy Micro RNA extraction kit (QIAGEN) according to the manufacturer’s instructions.

### Quantitative real-time PCR

RNA was extracted from the dissected PPN tissue using RNeasy mini kit (QIAGEN). cDNA was synthetized by using the SuperScript IV VILO Master Mix (Applied Biosystems) and preamplified for 14 cycles using TaqMan PreAmp Master Mix and pool of TaqMan Gene Expression Assays (Applied Biosystems). The resulting product was diluted and then used for PCR with the corresponding TaqMan Gene Expression Assay and TaqMan Fast Advanced Master Mix. Data were normalized to Hprt by the comparative CT method. The following TaqMan probes were used for PCR amplification of Hprt (Mm03024075_m1), Cacna1c (Mm01188822_m1), Cacna1d (Mm01209927_m1), Kcnj5 (Mm01175829_m1), Kcnj6 (Mm01215650_m1), Abcc8 (Mm00803450_m1) and Abcc9 (Mm00441638_m1).

### Analysis and statistics

Electrophysiology data were generally analyzed using Clampfit 10.3 software (Molecular Devices). For Fura-2 Ca^2+^ imaging combined with electrophysiological recording, Clampfit 10.3 and Igor Pro 5-8 (WaveMetrics) were used for analysis. Data are summarized using box plots showing median values, first and third quartiles, and range, unless otherwise specified. Statistical analysis was performed with GraphPad Prism 8 (GraphPad Software). Nonparametric tests (Mann-Whitney U test of significance or Wilcoxon signed rank test for unpaired or paired design experiments, respectively) were used, unless otherwise stated. Two-tailed tests were used unless the working hypothesis predicted a clear directionality to the change in outcome measure, in which case one-tailed tests were adopted. Normality was assessed using the Shapiro-Wilk test. Comparison of survival curves was performed with the log-rank (Mantel-Cox) test. Probability threshold for statistical significance was P<0.05.

## Acknowledgements

We wish to thank Sasha Ulrich for expert technical assistance. This study was supported by the JPB Foundation, NIH (NS 121174) and Aligning Science Across Parkinson’s [ASAP-020551] through the Michael J. Fox Foundation for Parkinson’s Research (MJFF).

